# Creating a highly directional anti-tumor immunity

**DOI:** 10.1101/169227

**Authors:** Vyacheslav A. Buts, Klavdiya P. Skibenko

## Abstract

A simple method of creating a highly directional anti-tumor immunity is proposed. It is shown that under the influence of ultrasound of small and medium intensity, it is possible to tear off antigens from the cell surface (surface antigens (SA)). It was found that the immunogenicity of these SAs is not less than the immunogenicity of intact cells. These results give a new opportunity for creating specific, acute-directed immunity against malignant tumors.

## 1. Introduction

There is now a revival of interest in research towards the development of antitumor vaccines (see, for example, [1, 2]). Special hopes in these studies are placed on new DNA and RNA technologies. In the most active areas of research, neoantigens are constructed in one way or another. In other cases, microorganisms are sought, whose antigenic properties are similar in their characteristics to neoantigens. All these areas of research are associated with great technical difficulties, with great financial costs. They are available to a small number of well-equipped laboratories. In addition, in all these cases it is difficult to create the antigens absolutely identical to neoantigens. Ideal would be the creation of only such antigens, according to which the immune system of the body would unambiguously identify the foreign tissue (surface antigens). Therefore, any thoughts, any studies in this direction are of interest. In the present work we report on the results, which, possibly, open such an opportunity.

Another interesting area in the fight against tumors is the use of ultrasound (US). Currently, the main efforts in these investigations were directed on the use of focused high intensity US (see, for example [3,4]). The effect of such US on the tumor leads to its destruction due to either the thermal effect or as a result of mechanical destruction of the tumor cells. In experiments with such US, in many cases, not only the tumor itself was destroyed, but also the suppression of metastases growth has occurred. The last result can be explained by the fact that the destroyed tumor cells secrete tumor antigens into the bloodstream, which lead to the activation of the immune system. In all these cases, it seems that the activation is nonspecific. Indeed, when the cells are destroyed, a huge amount of foreign antigens come out into the surrounding tissue. Some of them have the immunogenicity significantly larger the immunogenicity of the SA. Besides, there can appear the antigen competition. The author of the article [1] drew attention to the fact that modern researches in development of anti-tumor vaccines in many respects revive the well-known old trends. The same remark applies to US in the fight against tumors. Indeed, similar studies were conducted in the former Soviet Union in the late fifties of the last century (see [5-7]). In particular, experiments were conducted in which ultrasound of high intensity (up to 235 W/cm^2^) and a frequency of 1.5 MHz was directed to the tumor in a test tube. The cells were completely destroyed. Various fragments were extracted from the material obtained, which were used for immunize animals. In some cases, a positive result was observed with respect to the development of antitumor immunity. However, the results obtained were unstable. It was impossible to find stable unambiguous conditions for obtaining a positive result. Evidently, that ideal result would be to obtain not just fragments of destroyed tumor cells, but those antigens that are on the surface of tumor cells and only those, which can help the body’s immune system to recognize the tumor. The remaining antigens are harmful ballast. Currently, similar studies are being carried out at the Moscow University at the basic department of the Research Center “Kurchatov Institute” of the Faculty of General Physics and Molecular Electronics of the Physics Faculty.

It is clear that in all the cases listed above it is practically impossible to isolate the necessary fraction of antigens (surface antigens (SA)). However, it is known that the mechanisms of action of US on physical and biological objects are not restricted to their simple thermal action, their mechanical pressure on the interfaces of media, and cavitation. In fact, there is wide variety of other physical, chemical and biological mechanisms (see, for example, [8-10,14]). In particular, it is possible not to destroy tumor cells with US, but without destroying the cells to remove from their surface only surface antigens. For this reason, a series of experimental studies on the use of US with low and medium intensity to isolate SA was carried out. These antigens can be used for immunization. They are easily accessible to cells of the immune system, and their immunogenicity, as will be shown below, is not inferior to the immunogenicity of the intact cells. The results on this subject of some of the experimental studies carried out are contained in this paper.

## 2. Materials and methods

For the experiment a suspension of mutton erythrocytes with concentration 10^7^ cells in milliliter (10^7^ cell/ml) was prepared by three times flushing by physiological solution. This suspension was divided into two unequal parts. One part remained without modifications, and the second part was affected by ultrasound (US). The main experiments were performed with US of 0.5…1 W/cm^2^ intensity during 5 min and US frequency 2 MHz. Then both parts of suspension were centrifugalized during 10 minutes with ~10^3^ Rev/min. In such a way four components were obtained: E — erythrocytes without US acting, E_s_ — erythrocytes after US acting, L — a liquid obtained after a centrifuging of the erythrocytes without US acting, L_s_ — a liquid, obtained after a centrifuging of the erythrocytes after US acting.

Besides, some part of the suspension affected by US was centrifugalized one hour after US acting procedure. Thus, in addition to the enumerated components, two more components were obtained, which will be designated as: E_st_, L_st_. Finally, from these six components eight various mixtures were prepared: EL, EL_S_, E_S_L_S_, E_S_L, E_St_L, L_S_, L, L_St_. The number of erythrocytes in first five mixtures was the same, as well as in the control — 10^7^ cell/ml.

Each of these suspensions was used for inside peritoneal immunization of ten white non-breed mice. After immunization the whey of animal blood of each group were investigated on a agglutination reaction with erythrocytes that were irradiated with US and those not irradiated. It is necessary mark, that the US intensity used in the given series of experiments did not result in erythrocytes destruction.

The results of this experiment are represented in Table 1; the titers of agglutination reaction (AR) between mutton erythrocytes irradiated with US (E_S_) and unirradiated mutton erythrocytes (E) are shown together with the antibodies in the whey of blood of each mice group.

**Table 1.**
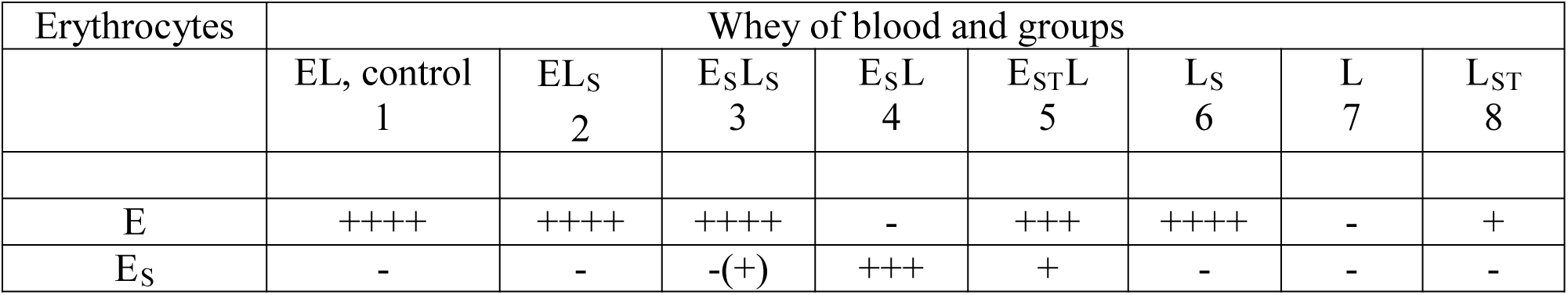

### 2.1 Discussion of results and some conclusions

It is easy to see using the information datas indicated in tab. 1 that all obtained results can be completely explained if to assume:

1. Under impact of low intensity US some cells lose the antigens from their surfaces and these antigens pass into a liquid surrounding the cells;
2. With time a portion of these antigens returns to surface of erythrocytes;
3. The structure of antigens and their immunogenicity do not change noticeably under the influence of low and medium US intensity.

The most important conclusion that follows from the obtained results can be formulated as: under impact of small and average intensity US, the tearing of antigens from surface of cells occurs; these antigens are easily separated from the cells in the course of centrifugation procedure. At that, the immunogenicity of the surface antigens (SA) is not less than the immunogenicity of cells which were exposed to US. This conclusion, to all appearance, does not depend on the type of cells.

This conclusion is supported by analysis of results of works [8,9]. Besides, we performed preliminary experiments on immunization of mice by liquid that was obtained after centrifuging of Erlix adenocarcinoma cells that were impacted by US. In the whey of blood of mice immunized with such liquid does always observe antibodies giving high titers of an agglutination reaction (AR) with Erlix adenocarcinoma cells not exposed to US. However, there was no such response when Erlix cells were irradiated by US.

The agglutination reaction (AR) of erythrocytes, exposed and not exposed to US, with antibodies in the whey of blood of the fourth group mice, gives evidence that already after 5 minutes of US exposure with intensity 0.5… 1 W/cm^2^ a vacation of the certain SA layer has occurred, and a new antigen layer became open. This new layer has greater resistance to US impact. To clear up the resistance of this layer, the immunogenicity of erythrocytes subjected to much higher US intensity (3 W/cm^2^) during longer time (20 min) was studied. It was found that the immunogenicity of these erythrocytes not too much differ from the immunogenicity of those erythrocytes that were exposed to low level of US during low time (I=0.5 W/cm^2^, t=5 min

It is important to note the almost complete absence of agglutination reaction (AR) between erythrocytes exposed to US and the antibodies in the blood of the third group of animals. Based on the components which took part in the immunization (E_S_L_S_) one could expect the presence of antibodies in both cases: with and without US.

The absence of the response on the US irradiated erythrocytes is explained as follows: during the resolution time of the administered suspension the majority of antigens “have occupied” the vacant places on the surface of erythrocytes and thus “have covered” the freed antigenic layer.

The confirmation of this conclusion can be the results of AR between mutton erythrocytes and antibodies of the blood whey of the mouses immunized by erythrocytes (E_ST_ L) and liquid (LST) irradiated by US followed seasoning process during one hour before centrifugation. The decrease of an AR titer in these groups (fifth and eighth) also is easily explained in the way: the SA, separated by US from the erythrocytes surface, have the tendency to return to the erythrocytes surface again.

## 3. Use US for creation of immunity against tumour Erlix adenocarcinoma cells (ACE)

It is natural to assume, that the mutton erythrocytes are not unique cells, from which surface the antigens can be separated by the use of US. Such US effect, probably, is characteristic for all cells, and it is possible to expect, that immune activity of the plucked antigens also will be high, as well as SA of the mutton erythrocytes. The results of experiments with tumour Erlix adenocarcinoma cells (ACE) confirm this supposition: compere results [11-12] and results [13].

In these experiments the ACE cells were irradiated by US with intensity 0.1…1 W/cm^2^ and frequency 2 MHz during 2…5 minutes. Then 0.3 milliliter of the mixture was hypodermicaly injected to outbred white mice for getting the solid form of tumor, and with 1 milliliter intraperitoneally in the case of ascites form. The animals were distributed into groups. The control group was injected with the ACE cells not US irradiated. The concentration of cells in the cell suspension was 10^7^ cell/ml in all groups.

Prior to inoculation, the percentage of viable cells was calculated by a supravital dyes method. The ability to vaccinate was determined by measuring number of nuclear cells in 1 ml of ascitic liquid extracted from abdominal cavity of the experimental animals on the tenth day after inoculation. The intensity of the growth of solid tumors was determined by analyzing their weight, also on the tenth day. The results of these experiments show a significant inhibition of tumor growth after exposure of the ACE cells to US irradiation. From Table 2 it is seen that the growth of solid tumors has slowed down more than 2.5 times. These results became the basis for further work.

**Table 2.**

| Suspension of ACE cells | Doze of irradiation. $\text{Wt/sm}^2$ | Number viable cells, % | Number of nuclear cells in 1 ml | Average weight of solid swellings (mlgr) | Average increasing of number ascitic cells on tenth day | Dispersion $\sigma$ |
| --- | --- | --- | --- | --- | --- | --- |
| Control | | 92 | $10^{11}$ | 220 | 35 | 65 |
| Experiment | 0.1...1 | 90 | $10^{11}$ | 82 | 22 | 10 |

After receiving these data, the scheme of experiments was complicated. The ACE cells, with and without US irradiation, were centrifuged. As the result, four fractions were obtained:

1. Supernatant of the ascite after US (L_S_);
2. The cells of ascite after US (Cs);
3. Supernatant of the ascite without US (L);
4. The cells of the ascite without US (C).

After centrifugation, the cells were washed twice with physiological solution. Then suspensions of cells were prepared, with cross replaced supernatants of the ascite: with and without US. Four “grades” of cell suspensions thus were obtained:

I - With cells not irradiated with US (control cells (CL));

II - Ascite, irradiated with US (C_S_L_S_);

III - The cells that were not US irradiated plus supernatant of the ascite that passed US irradiation (CL_S_);

IV - The cells of the ascite that was subjected by US, plus supernatant of the ascitet that was not subjected by US (C_S_L).

These cell suspensions were injected to four groups of outbred white mice by the method described above. It should be noted that the percentage of viable ACE cells, that passed US, do not differ from the control, amounting 95 …97%. The results of experiments are indicated in table 3.

**Table 3:**

| Groups | Cells viability, % | Number of nuclear cells in milliliter | Average arising of number ascite cells on tenth day | Average weight solid swelling on 15<sup>th</sup> day | Coefficient of variation - $V$, % | Relative error $\varepsilon$ % | Dispersion $\sigma$ |
| --- | --- | --- | --- | --- | --- | --- | --- |
| I CL | 95 | $4 \cdot 10^7$ | 36 | 682 | 39.1 | 30 | 270 |
| II $C_S L_S$ | 96 | $4 \cdot 10^7$ | 24 | 305.4 | 4.9 | 4 | 17.9 |
| III $CL_S$ | 94 | $4 \cdot 10^7$ | 29 | 225.3 | 8 | 6 | 14.8 |
| IV $C_S L$ | 97 | $4 \cdot 10^7$ | 21 | 139 | 15 | 12 | 23.7 |

For the processing of the results, the following statistical characteristics were used: dispersion σ, the coefficient of variation *V*, and relative experimental error *ε*. These values were calculated according to the formulas:

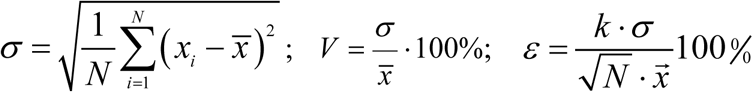

where N – is the number of mouse in experimental groups (≥ 10);

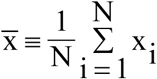 - mean tumor weight, *k*-numerical factor, in our case *k* ≈ 2.

Table 3 shows the results of processing of experimental data with the solid form tumors. As seen, the coefficient of variation V for all groups does not exceed 40 %. This indicates normal distribution of studied variables (weight of tumors) and the possibility of statistical processing of results. As follows from data of Table 3, tumor growth inhibition effect is observed in all experimental groups, and suppression of solid form of tumors is more essential than of ascitic form.

## 4. Conclusion

Thus, by exposure to low and medium intensity ultrasound, antigens can be disrupted from the cell surface. At that, the immunogenicity of these PAs is not inferior to the immunogenicity of the cells themselves. It means that the number of specific antibodies produced by the body on the introducing of PA is about the same as the number of antibodies of the same specificity that appear after the introduction of the intact cells into the body. If we talk about the development of an acute directed immune response, then an introduction of only PA into the body is certainly preferable in comparison with immunization with cells. Indeed, together with cells, a large number of other, in some cases “stronger” antigens are introduced. The presence of these antigens significantly weakens the immune response to surface antigens. It does occur, for example, due to the competition effect. But in fact the reaction to PA characterizes the effectiveness of the immune response to tumor cells. For the same reason, the use of high intensity US (I> 3 W / cm^2^) should be less effective because cavitation bubbles arising in the US wave fields lead to cells disruption. This consideration is now particularly important, since after successful experiments with intense US, attempts to govern the growth of malignant tumors with US generally follow along the path of increasing its intensity [3].

It should be noted that there have been attempts to use low and medium US to fight tumors. With these attempts, tumors in the body were exposed by US for a not too long time (up to 40 minutes). In such a case, the antigens, torn from the cell surface, as we saw above, can return to the surface of the cells and do not have time to manifest themselves. At the same time, it is known that under the influence of US of low and medium intensity, the nutrition of tissues is improved, i.e. stimulation of tumor growth occurs. These considerations easily explain the contradictory results of a direct short-term exposure of the tumor to ultrasound of medium intensity. From here follows also the possibility of effective antitumor activity of ultrasound of medium and low intensity with prolonged or even permanent exposure of the tumor. However, the most valid and effective at the moment schema of using US to combat with already formed tumor is as follows. The tumor or tumor cells are removed from the body. Then a suspension of SA is prepared from these cells with the help of US of low or medium intensity. This SA is injected into the body to produce acute-directed immunity, followed by destruction of the remaining cells and metastases. Of course, infinite complications and variations of this scheme are possible. For example, one can additionally use the adjuvants or use additional nonspecific stimulation of the immune system, etc.

There is no doubt that the surface antigens from microbes can be produced by appropriate selection of the US intensity and frequency. The presence of such antigens provides an ideal material for vaccination. Note that surface antigens can also be used for the early diagnosis of some tumors. The lack of an immune response, even with a large number of malignant tumor cells, can be in particular due to the presence of an increased Coulomb potential of tumor cells. This potential makes it difficult to collide and recognize these cells with immunocompetent cells [10]. With the help of US of small and medium intensity, the Coulomb potential of cells can be reduced. In our preliminary experiments we observed a noticeable decreasing in the external electric field of the mobility of the sounded sheep erythrocytes and ACE cells compared to the mobility of non-sound ones, although the quantitative regularity of the change of the potential and its dynamics were not studied. Reduction of the Coulomb potential of tumor cells, as mentioned above, allows to strengthen the immune response to these cells. At the same time, reduction of the potential of normal cells is a negative effect, as, for example, for blood cells, it can lead to their aggregation and, as a consequence, to the occurrence of thrombi.

The authors are grateful to A.A. Tsutsaeva for numerous discussions of the results and for useful advices. The authors also thank V.S. Voitzeny for discussion and for editing English text.

